# Relevance study of vector competence and insecticide resistance in *Aedes aegypti* laboratory lines

**DOI:** 10.1101/2022.02.12.480187

**Authors:** Lanjiao Wang, Albin Fontaine, Pascal Gaborit, Amandine Guidez, Jean Issaly, Romain Girod, Mirdad Kazanji, Dominique Rousset, Marco Vignuzzi, Yanouk Epelboin, Isabelle Dusfour

**Affiliations:** Vectopôle Amazonien Emile Abonnenc, Unité de contrôle et adaptation des vecteurs, Institut Pasteur de la Guyane, 23 avenue Pasteur, 97306 Cayenne cedex, French Guiana, France; Unité de Parasitologie et Entomologie, Département des Maladies Infectieuses, Institut de Recherche Biomédicale des Armées, 19-21 Boulevard Jean Moulin, 13005 Marseille, France; Aix Marseille Université, IRD, AP-HM, SSA, UMR Vecteurs – Infections Tropicales et Méditerranéennes (VITROME), IHU – Méditerranée Infection, 19-21 bd Jean Moulin, 13385 Marseille, cedex 5, France; Institut Pasteur de la Guyane, 23 avenue Pasteur, 97306 Cayenne cedex, French Guiana, France; Laboratoire de Virologie, Institut Pasteur de la Guyane, 23 avenue Pasteur, 97306 Cayenne cedex, French Guiana, France; Unité des Populations Virales et Pathogénèse, Institut Pasteur, 28 rue du Dr Roux, 75724 Paris cedex 15, France

## Abstract

The urban mosquito species *Aedes aegypti* is the main vector of arboviruses worldwide. Mosquito control with insecticides is the most prevalent method for preventing transmission in the absence of effective vaccines and available treatments; however, the extensive use of insecticides has led to the development of resistance in mosquito populations throughout the world, and the number of epidemics caused by arboviruses has increased.

Three mosquito lines with different resistance profiles to deltamethrin were isolated in French Guiana, including one with the I1016 knock-down resistant allele. Significant differences were observed in the cumulative proportion of mosquitoes with a disseminated chikungunya virus infection over time. In addition, certain genes (*CYP6BB2, CYP6N12, GST2, trypsin*) were variably overexpressed in the midgut at 7 days after an infectious blood meal in these three lines. Therefore, detoxification enzymes and *kdr* mutations may contribute to an enhanced midgut barrier and reduced dissemination rate.

Our work shows that vector competence for chikungunya virus varied between *Ae. aegypti* laboratory lines with different deltamethrin-resistance profiles. More accurate verification of the functional association between insecticide resistance and vector competence remains to be demonstrated.

**Importance:** Three *Ae. aegypti* lines, isolated from the same collection site, underwent different insecticide selection pressures against deltamethrin under laboratory conditions. As a result, they developed different resistant profiles. In this study, when these lines were fed an artificial infectious blood meal containing chikungunya virus, all three lines including the reference strain showed a high infection rate. There was no statistical difference in infection rate found; however, the dissemination rate of the virus from midgut to head were significantly different. A higher resistance level detected by the WHO test was correlated with a lower viral dissemination rate for each strain. This study presented evidence that the insecticide selection pressure or the existence of insecticide resistance could lead to differences in viral dissemination or even transmission in mosquito populations. We hope that our study can give more insights into understanding the roles of mosquito insecticide resistance on viral transmission.

## Introduction

*Aedes aegypti* (Linnaeus, 1762) is a major vector of arboviruses worldwide and, as such, has been the target of insecticide control strategies for decades (Carvalho and Moreira, 2017). Understanding the interactions involved in vector competence and deciphering the origins of insecticide resistance have been the focus of many research topics, aiming to improve the control of this vector and disrupt virus transmission.

Taken together the vector competence studies of *Ae. aegypti* for dengue virus, Zika virus, chikungunya virus and yellow fever virus from around the world, vector competence varied when different mosquito populations and virus strains were used (Souza-Neto, Powell and Bonizzoni, 2019). Other factors such as the presence of symbiont microbiota and non-pathogenic mosquito-specific viruses may contribute to the impact on the susceptibility of mosquitoes to viral infection as well (Souza-Neto, Powell and Bonizzoni, 2019). The virus infection initiates in the midgut and then reaches the salivary glands from where the virus can be expectorated to the saliva and transmitted to another vertebrate host. Therefore, midgut barrier and salivary gland barrier especially the former is the key point for the assessment of vector competence of mosquitoes. For instance, it was shown that there was no danger of *Ae. aegypti* transmitting the live attenuated yellow fever 17D vaccine (Whitman, 1939), because during natural infection, the block for yellow fever 17D replication occurs at the basal membrane of the midgut (Danet *et al*., 2019).

It was shown that a major component of the basal lamina of the extracellular matrix surrounding the midgut will be temporally degraded by collagen IV during an arbovirus infection. It can allow the midgut barrier to become more permissive for the virus and resulting in a significantly increased chance of escape from the midgut (Dong *et al*., 2017). The ultrastructural analysis of the *Ae. aegypti* midgut also revealed the interactions between bloodmeal digestion and the quick viral dissemination within 48h post blood feeding (Kantor *et al*., 2018). Additionally, based on the comparative differential expression analysis of midguts from saline meal-feeding mosquitoes and protein meal-feeding mosquitoes, midgut-expressed genes coding for trypsins, metalloproteinases, and serine-type endopeptidases were not involved in blood or protein digestion, but could contribute to the midgut escape for CHIKV in *Ae. aegypti* (Dong, Behura and Franz, 2017).

To date, *Ae. aegypti* populations all around the world have developed insecticide resistance, especially to pyrethroids and organophosphates (Moyes *et al*., 2017). Resistance to deltamethrin has been the most widely monitored and its mechanisms studied (Dusfour *et al*., 2015). Researches indicate that resistance may be conferred by target-site modification due to non-synonymous mutations in the voltage-gated sodium channel sequence, causing knock-down resistance (*kdr*). Changes in the amino-acid sequence of the target prevent the molecule from correctly binding and reducing its toxic action. (Dong *et al*., 2014; Du *et al*., 2016). Currently, at least 10 mutations on the voltage-gated channel sequence have been associated with pyrethroid resistance in *Ae. aegypti* populations worldwide (Du *et al*., 2016; Granada *et al*., 2018). The presence of 1016I and 1534C mutations in *Ae. aegypti* are reported to be strongly related to deltamethrin and permethrin resistance in the Americas (Saavedra-Rodriguez *et al*., 2007; Dusfour *et al*., 2015), and double mutants have an enhanced effect on resistance to deltamethrin (Brito *et al*., 2013).

Metabolic resistance is another major mechanism of insecticide resistance. It is caused by an increase in detoxification enzymes or the production of more efficient isoforms, of which the cytochrome P450 (CYP450), glutathione-*S*-transferase (GST) and carboxy/cholinesterase gene (CCE) family have mainly been studied. These changes improve the degradation, sequestration or excretion of toxic molecules (Hemingway *et al*., 2004). Resistance is related to the geographical origin of the populations that were investigated and probably to past insecticide practices (Faucon *et al*., 2015).

In studies focusing on the interactions between vector competence and insecticide resistance, evidence of interactions between insecticide resistance and the incidence of parasite infections has been found for *An. gambiae–Plasmodium falciparum* and *Cx. quinquefasciatus–Wuchereria bancrofti* (McCarroll and Hemingway, 2002; Alout *et al*., 2013). In addition, Atyame et al. presented evidence that insecticide resistance could enhance the vector competence of *Cx. quinquefasciatus* for West Nile virus (Atyame *et al*., 2019). More recently, two researches focused on influence of vector competence of pyrethroid resistant *Ae. aegypti* for Zika virus, the results were similar between the resistant group and susceptible group (Dos Santos *et al*., 2020; Parker-Crockett *et al*., 2021), however a reduced vector competence was found in resistant *Aedes albopictus* for dengue virus (serotype 2) (Deng *et al*., 2021).

In this study, we set up three isofemale lines isolated from a population in French Guiana where deltamethrin resistance in *Ae. aegypti* is extremely high (>1000 times) and associated with the presence of 1016I, CCE and CYP450 gene over-expression including *CYP6BB2, CYP6M11, CYP6N12, CYP9J9* and *CYP9J10* (Dusfour *et al*., 2015; Faucon *et al*., 2015). In addition, field populations of *Ae. aegypti* mosquitoes from French Guiana has shown a high susceptibility to CHIKV infection (Girod *et al*., 2011). Herein, in this study we will investigate the vector competence of CHIKV amongst three French Guiana isolating laboratory strains, aiming to learn more about the insecticide resistance, vector competence and potential interplay in *Aedes* mosquitoes.

## Methods

### Ethics statement

The experiments were authorized by agreement number B973-02-01 delivered by the Préfecture of French Guiana, which was renewed on 6 June 2015. The protocol for use of mice and rabbits was approved by the Committee for Ethics in Animal Experimentation of the Institut Pasteur (No. 89), report number 2015–0010 issued on 18 May 2015. All experiments were performed in accordance with relevant guidelines and regulations.

### Mosquitoes

Four laboratory strains of *Ae. aegypti* were used. Three of 60 lines produced by isolating F1 females from a natural population collected at Ile Royale, French Guiana (5.287° N, 52.590° W), were selected for their contrasting deltamethrin resistance profiles. Siblings were allowed to mate in subsequent generations. IR03 and IR05 were maintained under deltamethrin selective pressure by eliminating 75% of larvae every two generations up to the 12th generation, while IR13 was maintained in the same environmental conditions, but without any deltamethrin selective pressure up to 5th generation. The New Orleans (NO) strain, which is susceptible to all insecticides, was used as the reference in all experiments. Mosquitoes were reared under controlled environmental conditions of 12 h:12 h photoperiod, 80% ± 10% relative humidity and 28 ± 2 °C. Mosquitoes were fed on blood taken from anaesthetized mice. More details about the production and maintenance of the strain were described previously (Epelboin *et al*., 2021).

### Insecticide resistance detection

#### Bioassays: time and dose–response tests

Three lines, IR03, IR05 and IR13, were isolated from a single female in order to limit genetic variation and produce contrasting deltamethrin resistance phenotypes and mechanisms, in order to study the relations among vector competence phenotypes. Three to five days old females from the three lines (IR03, IR05 and IR13) and the NO strain were exposed to doses of 0.001–0.4% deltamethrin that cause 0–100% mortality. Technical-grade deltamethrin (Sigma Aldrich, Germany) dissolved in acetone and mixed with silicon oil was applied on Whatman paper according to the World Health Organization (WHO) recommended protocol (https://apps.who.int/iris/handle/10665/69296). For each line, we recorded: (i) the cumulative number of knocked-down (KD) mosquitoes exposed to 0.06% deltamethrin (i.e. the diagnostic dose for resistance monitoring) every 3 min for the first hour, and (ii) the mortality rate after 24 h (% 24-h M) at each dose. The theoretical time or dose that knocked down or killed 50% of the population (KDT50 and LD50, respectively) was obtained by probit regression with BioRssay in R statistical environment version 3.2.0. The resistance ratio, which is the LD50 of the tested population divided by the LD50 of the reference strain (RR50), was used to evaluate the level of resistance.

#### Knock-down resistance (kdr) genotyping

With DNA extracted from the midguts of all individuals, 103 of IR03, 159 of IR05, 136 of IR13 and 234 of NO adults were genotyped in discrimination allele assays to localize resistant alleles in positions 1016 and 1534 of the sodium channel voltage-dependent gene in a StepOnePlusReal-time PCR system (Life Technologies, Gaithersburg, MD, USA). The detailed protocol is described in Table S2.

#### Metabolic resistance related gene expression

A subset of individuals was selected 7 days post-infection with chikungunya, when 100% of the midguts and about half the heads were positive in each line, and the expression of genes associated with resistance to deltamethrin was compared in midguts with positive heads and those with negative heads. We chose 27, 30 and 30 positive midguts of IR03, IR 05 and IR13, respectively, of which 18, 16 and 18 had disseminated heads and 9, 14 and 12 had non-disseminated heads, respectively. RNA was extracted from midguts with an RNA kit from Qiagen, according to the manufacturer’s instructions. The concentration and quality of total RNA were determined in a spectrophotometer (NanoDrop 2000c Thermo Scientific). The total RNA yield was standardized at 60 ng/μL for all samples and then treated with DNAse (Carlsbad, CA, USA). Complementary DNA (cDNA) strands were synthesized with SuperScript™ III Reverse Transcriptase kit Invitrogen (Carlsbad, CA, USA), according to the manufacturer’s protocol. RpL8 was used as the housekeeping gene for CYP6BB2, CYP6N12, GST2 and trypsin. Quantitative RT-PCR was performed according to the protocol of Power SybrGreen PCR Master Mix (Applied Biosystems, Foster City, CA, USA). The detailed protocol is described in Table S3.

#### Vector competence assay

All experiments involving mosquito oral infections and virus manipulation were performed in a BioSafety Level-3 Laboratory following French regulation.

Chikungunya virus (CHIKV): Mosquitoes were infected orally with a CHIKV infectious clone produced by the Viral Populations and Pathogenesis Unit, Institut Pasteur, Paris, at a titre of 2.5 × 10^7^ PFU/mL from four PCR amplicons of patient serum obtained during the outbreak of chikungunya in the Caribbean islands in 2013. This clone was found to replicate in VERO cells like an Asian lineage of CHIKV (NC-2011) in baby hamster kidney cells. It is used to study pathogenesis and viral evolution (Stapleford *et al*., 2016).

#### Oral infection

Five-day-old females from the three selected lines and the NO strain were exposed to an infected blood meal via an artificial membrane feeding system (Discovery Workshops, Lancashire, United Kingdom) 24 h after being deprived of sucrose solution. The infectious blood meal was composed of fresh rabbit blood (volume proportion: 96%) and a final viral suspension of 10^6^ PFU/mL (volume proportion: 4%, with an initial titre of 2.5 × 10^7^ PFU/mL. 30 μL of ATP was added as a phago-stimulant at 5 mmol/L. After 30 min of exposure, the females were cold-anaesthetized, and fully engorged mosquitoes were maintained for up to 10 or 14 days with a constant supply of 10% sugar solution.

#### Sample collection

The vector competence of all strains was scored in 30 samples 3, 5, 7 and 10 days post-infection (DPI) for all strains according to three phenotypes: (i) midgut infection, as determined by the presence of virus in the midgut; (ii) virus dissemination from the infected midgut, as determined by the presence of virus in the head; and (iii) virus transmission, as determined by the presence of virus in the saliva of a mosquito with a disseminated infection. An extra 14 DPI time was used only for the NO strain, which had the highest engorgement rate and sample size. First, mosquitoes were cold-anaesthetized, their legs and wings were removed, and the proboscis was inserted into a filter tip ART (Molecular BioProducts, USA) with 20 μL of fetal bovine serum (FBS; Glico by Life Technologies). Saliva was collected for 30 min and stocked in tubes containing 130 μL of cell culture medium (L-15 Medium, Sigma). One limitation of this technique, which is commonly used for vector competence, is that the volume of saliva delivered by females cannot be estimated. The heads were then cut off and placed into tubes containing 500 μL of 20% FBS cell culture medium. Finally, the midguts were dissected and placed in a tube with 150 μL of 10% FBS cell culture medium. All samples were preserved at –80 °C.

#### Viral detection by real-time quantitative reverse transcription PCR (qRT-PCR)

The head was homogenized by grinding in microbeads for 20 s at 40 Hz (Biospec, USA). Then, 150 μL of supernatant were removed for RNA extraction with the QIAamp Viral RNA Kit (Qiagen, Germany). Midguts and saliva were extracted directly with the same kit after complete digestion in lysis buffer for 30 min at room temperature (about 25 °C). Finally, qRT-PCR was performed on a Stepone software v2.3 system instrument with SuperScript® III Platinum^®^ One-Step qRT-PCR Kit w/ROX (Invitrogen, CA, USA). The detailed protocol is shown in Table S1. Quantification was performed using standard curve, the limitation of detection is 100 genome copies.

#### Vector competence analyses

To analyse virus dynamics further, a logistic model with three parameters was used to describe the cumulative change in the proportion of mosquitos with a virus infection or dissemination over time post-exposure for each mosquito line by least-squares non-linear regression with the drm function in the drc R package (Cedergreen, Ritz and Streibig, 2005). Virus dissemination was scored only in mosquitoes with an infected midgut. The three-parameters logistic function (L.3 equation in the drm function with a self-starter) is given by the formula:

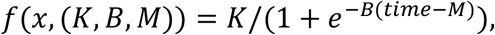

Where *K* represents the maximum prevalence of mosquitoes with a systemic infection (the saturation level), *M* the time at which the proportion of mosquitoes with a systemic infection equals 50% of the saturation level *K* (a proxy for EIP_50_) and *B* the growth rate around *M* (rate of change per time unit during the exponential phase). The growth rate (*B*) was further transformed into Δt, which corresponds to the rise time of the exponential function (time taken for a virus to rise from 10% to 90% of its saturation level), from the formula: log (81)/*B*. The logistic curve was forced to start at the origin for all tested lines. The sample size at each time was used as a weight argument.

The dynamics of lines were compared in an extra sum-of-square F test, which is based on the difference between the residual sum-of-squares of two models adjusted for the difference in the number of degrees of freedom: (i) a restricted model fitted to the combined sets of data and (ii) an unrestricted model fitted to each data set separately. The restricted model corresponds to a single sigmoid curve with three parameters estimated from the entire data set, whereas the unrestricted model corresponds to two distinct curves with three parameters estimated from each data set. The null hypothesis of the extra sum-of-square F test is that the unrestricted model does not fit the data better than the restricted model. In essence, if two separate curves fit the data better than a single curve, the null hypothesis is rejected, and the two curves are considered distinct. The F test quantifies improvement in the unrestricted model over the restricted one and generates a *P* value based on the F distribution and the number of degrees of freedom.

The same test was used to compare single parameters between two curves. In this test, the restricted model is defined as having the same value for one parameter in both data sets and estimates the two other parameters separately. The null hypothesis is that both data sets have the same shared parameter. If two separate curves fit the data better than the same curves constrained to share one parameter, the null hypothesis is rejected, and the shared parameter is considered distinct. The *P* value is derived from the F test on the basis of the F distribution and the number of degrees of freedom.

Virus title differences among lines were analysed in an ANOVA that included the effect of time post-virus exposure. All statistical analyses were performed in the R statistical environment version 3.2.0, and charts were generated with the ggplot2 package.

## Results

### Contrasted resistance profiles to deltamethrin

In the IR03 and IR13 *Ae. aegypti* laboratory lines, 100% of females were knocked down 1 h after exposure to deltamethrin at the diagnostic dose of 0.06%, with a KDT_50_ equivalent to that of the NO reference strain (Table 1). In IR05, 40% of mosquitoes were knocked down after 1 h, with a KDT_50_ extending over 1 h (Table 1). After 24 h of observation, 96% of IR13 females died at the dose of 0.06%, IR03 at 30.4% and IR05 at 26.4%. The RR_50_ was 6.42, 69.5 and 51.5 for IR13, IR05 and IR03, respectively (Table 1). The profile of IR13 was close to that of NO, with full susceptibility to the KD effect and low resistance. IR05 showed a high KD effect and resistance, while IR03 showed full susceptibility to the KD effect and resistance to deltamethrin. The presence of a KD effect and loss of susceptibility suggested the presence of metabolic resistance only in IR03 and at a lower level in IR13.

**Table 1.** Results of bioassays.

| Strain | % KD 1 h | KDT<sub>50</sub> [95% CI]<br>(min) | % 24 h M | RR<sub>50</sub> [95% CI] |
| --- | --- | --- | --- | --- |
| IR03 | 100% | 6.7 (6.1-7.3) | 30.4% (± 5.9) | 51.5 (37.3-71.2) |
| IR05 | 40% (± 4.8) | > 60 | 26.4% (± 4.0) | 69.5 (50.8- 95.2) |
| IR13 | 100% | 5.2 (4.6-5.8) | 96% (± 2.3) | 6.42 (4.6-9.0) |
| NO | 100% | 5.2 (4.7-5.6) | 100% | 1 |

% KD 1 h: percentage of mosquitoes knocked-down after 1 h of exposure to paper impregnated with 0.06% deltamethrin; KDT<sub>50</sub>: theoretical time to knock down 50% of tested females exposed to 0.06% deltamethrin; % 24 h M: percentage mortality of females after 24 h of observation; RR<sub>50</sub>: resistance ratio and its 95% confidence interval

### High infection rates in all lines

In total, 632 mosquitoes of three resistant lines and one reference strain were successfully analysed. All the strains were highly susceptible to the oral infection of CHIKV, with infection rates in midguts of > 90% as early as at 3 days post-infection (DPI). All mosquitoes of the IR13 line were infected (S1 Fig. A). The midgut infection rates of the three lines were significantly different but were not significantly influenced by time post-exposure, regarding of further analyses of infection dynamics in the midgut (logistic regression analysis of deviance: *P*=0.003 for the mosquito line effect, *P*=0.97 for the effect of time post-exposure and *P*=0.56 for their interaction). Pairwise comparisons among the lines, except for time, showed significant differences between the low (IR13) and the high (IR05) resistance lines (*P*=0.031).

Virus load in infected midguts was determined at each time post-exposure in all lines. The load also increased significantly (ANOVA, *P*=0.0001) with time post-exposure for all lines.

### Significant differences in dissemination saturation levels

Mosquitoes in each line with an infected midgut were scored for disseminated infection over time, and the dynamics of disseminated infection was fitted to a logistic equation (sigmoid shape) to derive the three parameters that best described the virus dissemination kinetics in each mosquito line (Fig. 1). Our model was not optimal for the IR13 line, as an additional time after 10 days’ post-virus exposure would have properly captured the dissemination plateau (saturation level). Therefore, the disseminated infection saturation level for this line is probably underestimated.

**Fig 1.**
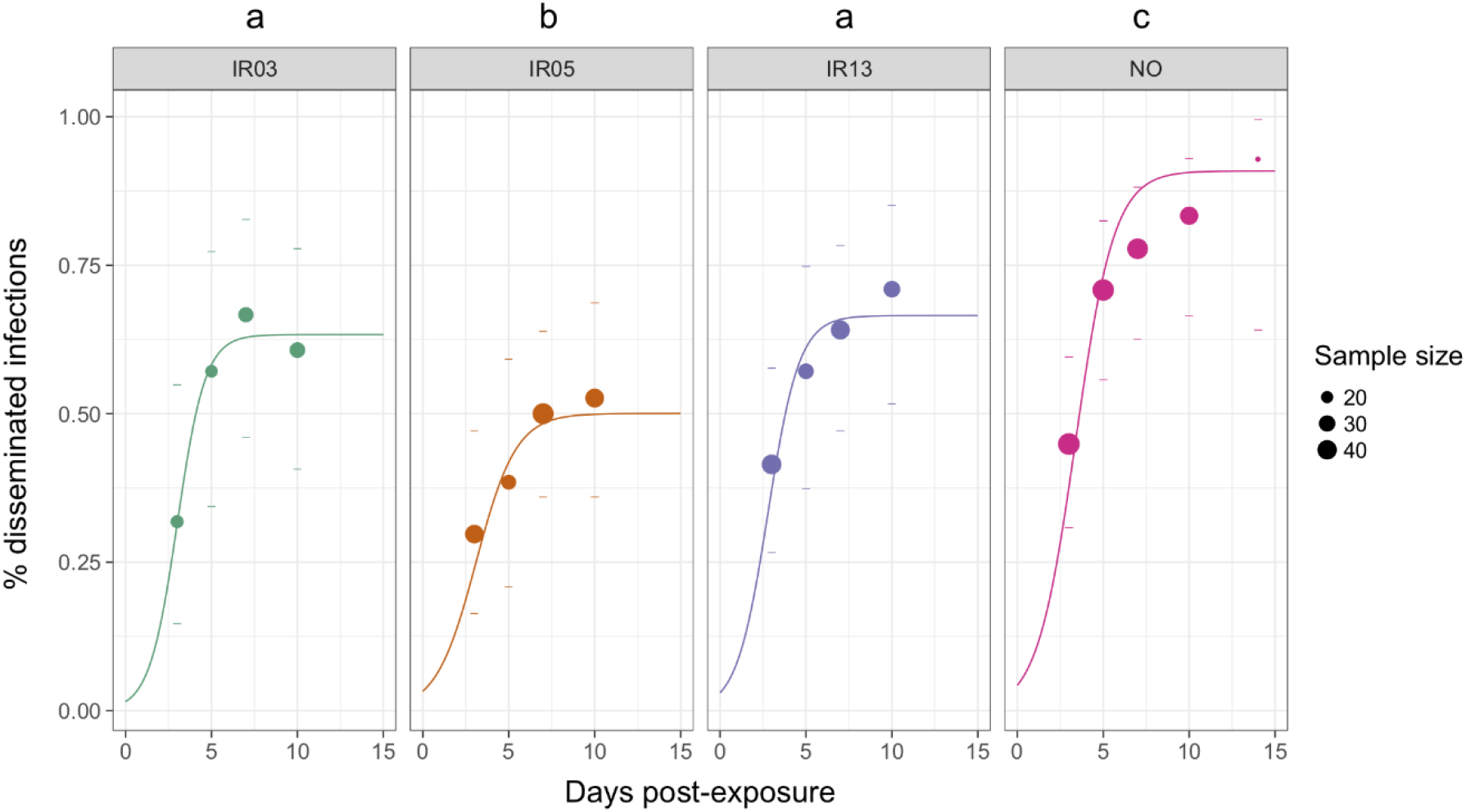
Cumulative proportion of mosquitoes with viral dissemination from the infected midgut over time for each mosquito line and the reference NO strain.

No significant difference was found between the lines in time-dependent parameters, i.e. the time at which the proportion of mosquitoes with a disseminated infection is 50% of the saturation level (which corresponds graphically to the point of symmetry of the sigmoid) and the time required to increase from 10% to 90% of the saturation level (Δt, the rising time of the sigmoid). All mosquito lines reached their dissemination saturation level (K) at about 7 days, and the time to reach 50% of the disseminated infection proportions (M) was about 3 days for all lines. Significant differences were observed between lines in saturation level (ANOVA, IR05 and IR03, *P*=0.04; IR05 and IR13, *P*=0.04), which corresponds to the maximum proportion of disseminated infection that can be attained in the line. IR05 showed 50% virus dissemination, IR03 63% and IR13 an underestimated 66%. The value for the reference NO strain was 91% of dissemination (Table 2).

**Table 2.** Estimated parameters of the dynamics of disseminated infections.

| Mosquito<br>line | K | B | Δt | M |
| --- | --- | --- | --- | --- |
| IR03 | 0.63 | 1.23 | 3.6 | 3.0 |
| IR05 | 0.50 | 0.86 | 5.1 | 3.1 |
| IR13 | 0.67 | 1.10 | 4.0 | 2.8 |
| NO | 0.91 | 0.89 | 5.0 | 3.4 |

K: maximum prevalence of mosquitoes with a disseminated infection (saturation level); M: time to reach 50% of the saturation level K; B: the growth rate around M; Δt: the rise time of the exponential function (time for a virus to rise from 10% to 90% of its saturation level).

Regarding the viral load in midgut, the threshold was around 10^5 genome copies / midgut which could lead to a successful crossing escape from midgut to reach other organs such as head (Fig. S3). The viral loads in individual heads were significantly influenced by the time post-exposure (*P*=4.93 × 10^−6^) and by the mosquito line (*P* < 0.02) (Fig. S2). Differences among the mosquito lines found only for the IR13 and IR05 pair (*P*=0.009 adjusted for multiplicity of tests).

### No evident difference found regarding transmission rate

The transmission rate is the percentage of mosquitoes with positive saliva among those with positive heads, and the transmission efficiency rate is the percentage of mosquitoes with positive saliva among all mosquitoes tested. Neither could be modelled, because, in certain cases, < 10 positive saliva samples were obtained (Table S4). As dissemination was stabilized from 7 DPI and sample sizes were highest at 7 and 10 DPI, the transmission rates ranged from 48% to 85% and transmission efficiency from 23% to 55%. Chi-square comparison did not show any difference between the lines at both 7 and 10 DPI.

### Resistant mutation at the 1534 position is inversely associated with the dissemination saturation level

The frequency of resistant allele 1016I accorded with *kd* phenotypes and our knowledge of *kd* resistance mechanisms in French Guiana populations. IR05 was the only line with a high frequency of the 1016I resistant allele (0.804) and the three genotypes with a predominance of homozygote resistance (64.1%) (Fig. 2). The 1534 locus displayed various proportions of the 1534C resistant allele and all three genotypes, depending on the lines. IR05 had a frequency of 0.814 for the resistant allele, bringing the proportion of double homozygote resistant (I1016I/C1534C) to 63.5% and that of full heterozygotes up to 30.8% in this line (Fig. 2). The frequencies of other genotypes were null or negligible (< 3%). The frequency of 1534C in IR03 was 0.456.

**Fig 2.**
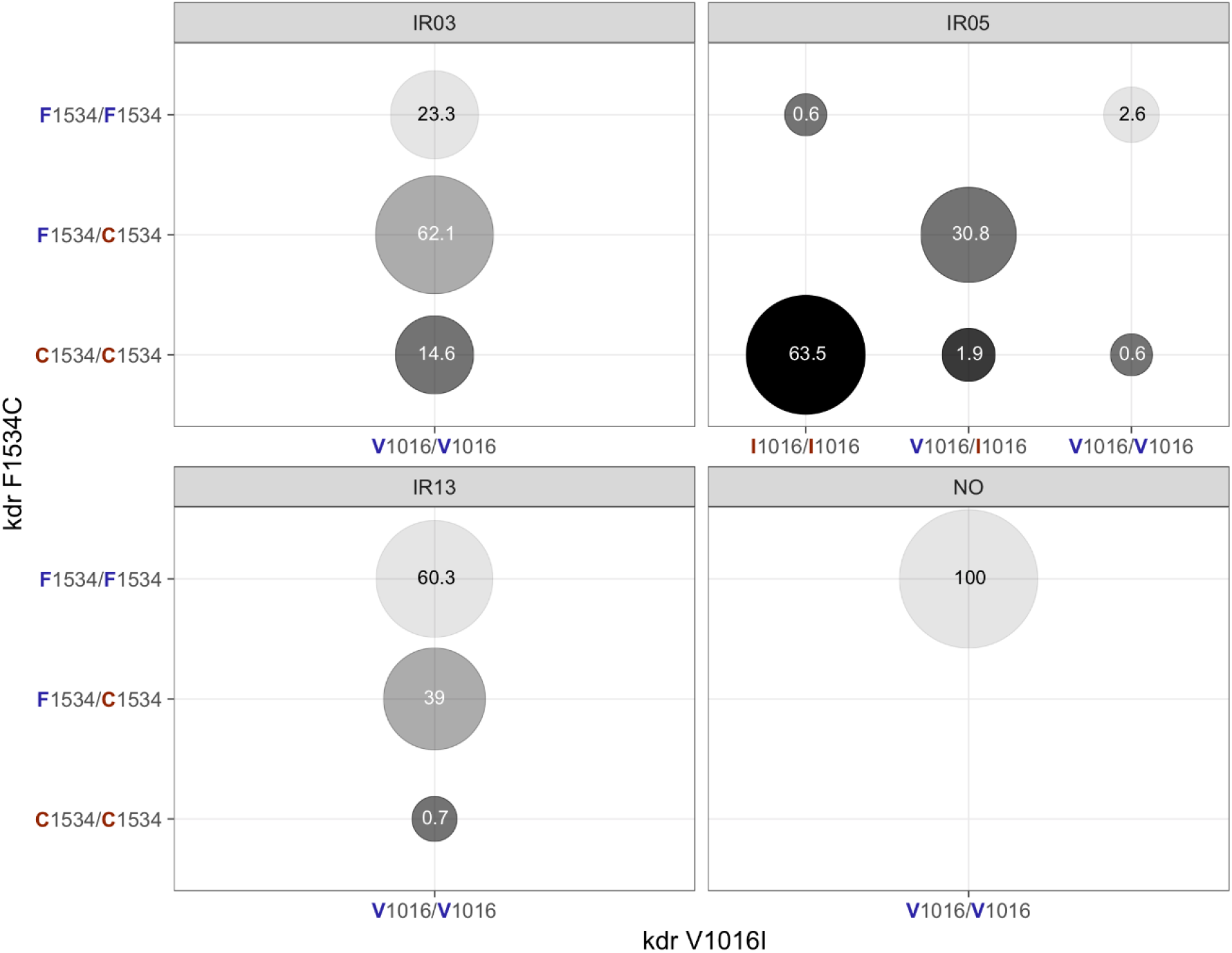
Genotype proportions of mutations V1016I and F1534C involved in deltamethrin resistance in each mosquito tested line and the reference NO strain. Proportions (%) of mosquitoes with each combination of alleles for these two mutations are represented in circles. Susceptible alleles are shown in blue letters and resistant ones in red. The colours of the circles refer to the number of resistant alleles in each genotype, becoming darker with the number of resistant alleles.

The proportion of heterozygotes was 62.1%, and those of susceptible and resistant homozygotes genotypes were 23.3% and 14.6%, respectively. IR13 had the lowest frequency of resistant 1534C (0.202) and the highest proportion of homozygote susceptible genotypes (60.3%) (Fig. 2). The proportion of heterozygotes reached 39%, while that of the resistant homozygote genotype was negligible (0.7%). As expected, the susceptible alleles 1016V and 1534F were fixed in the NO reference strain.

When the dissemination saturation level is taken into consideration with the proportion of 1534C by mosquito line, an interesting inverse relation appears (Fig. 3). No difference in the virus loads in midguts or heads was observed among the lines with different *kdr* genotypes (S1 Fig B).

**Fig 3.**
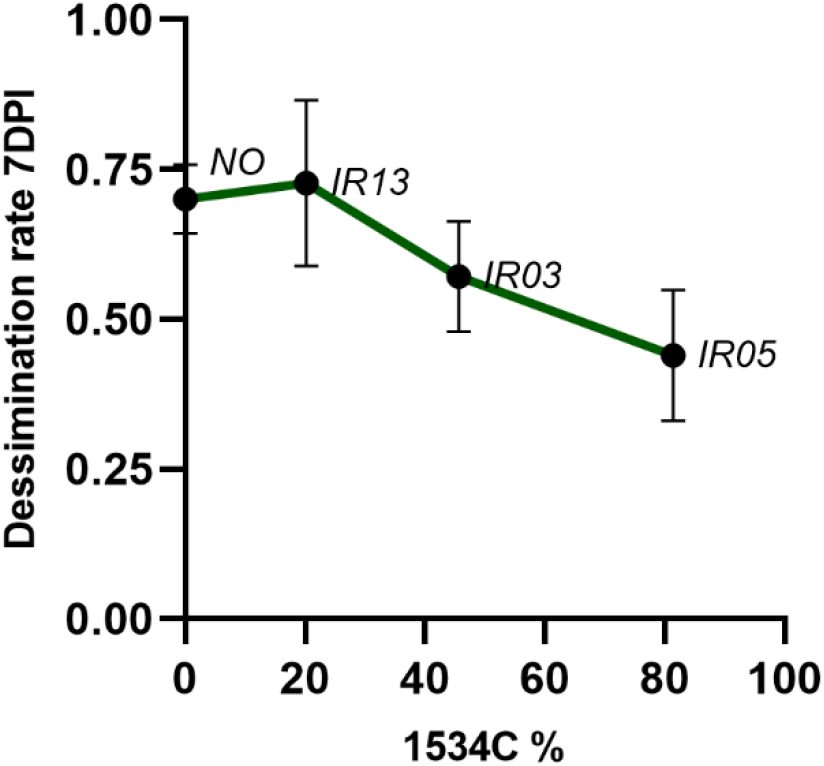
Relation between dissemination saturation level and proportion of resistant alleles 1534C.

### Relative expression level of detoxification enzyme varied in the midgut

A sub-sample of 87 midguts tested 7 DPI had a frequency of 1016I of 0.455 and a frequency of 1534C of 0.47. In the V1016I mutation, the proportion of heterozygotes was 12.6%, and those of susceptible and resistant homozygotes genotypes were 67.8% and 19.6%, respectively. In the F1534C mutation, the proportions were 46.0% heterozygote, 30.0% susceptible and 24.1% resistant homozygote genotypes. RpL8 was used as the housekeeping gene to quantify the expression level of CYP6BB2, CYP6N12, GST2 and trypsin. There is no significant difference fund between the group of midguts with positive heads and the group of midguts with negative heads.

Relative expression level was shown by Fig 4, CYP6BB2 was more expressed in IR03 and IR05 than in IR13 (Mann-Whitney, IR03 and IRI3, *P*< 0.0001; IR05 and IR13, *P*<0.0001, Fig. 4A). Regarding the expression level of CYP6N12, there is no significant differences when compare IR03 with IR05 or IR13, however it was more expressed in IR05 than in IR13 (Mann-Whitney, *P*=0.0016, Fig. 4B). Looking at the expression of GST, similarly, there is no significant differences found between IR03 and IR05, but they both have a higher expression ratio than IR13 (Mann-Whitney, IR03 and IRI3, *P*=0.0005; IR05 and IR13, *P*<0.0001, Fig.4C). Finally, more significant variation was found for the results of trypsin, only in IR03, it was overexpressed, it was expressed negatively in both IR05 and IR13, the difference was significant among three lines (Mann-Whitney, IR03 and IRI3, *P*=0.0001; IR05 and IR13, *P*<0.0307; IR05 and IR03, *P*=0.0038, Fig. 4D).

**Fig 4.**
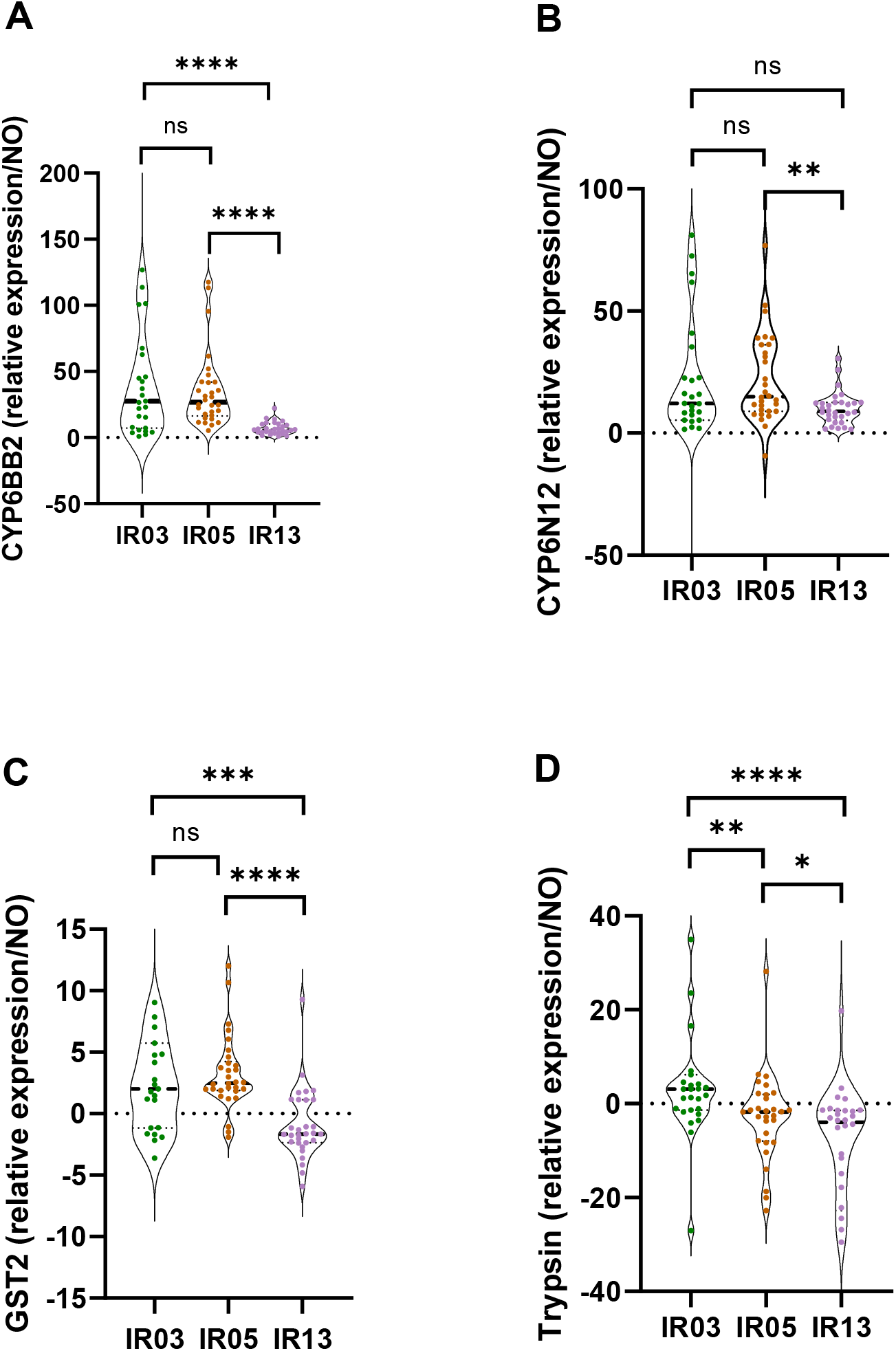
Expression ratio of *CYP6BB2, CYP6N12, GST2* and *trypsin* standardized by the housekeeping gene *RpL8* in IR03, IR05 and IR13 compared with NO stain.

## Discussion

The three isofemale lines originated from the same breeding site but were raised under different levels of deltamethrin-selecting pressures. Distinct phenotypes and target site mutation profiles were detected among them. IR05 was highly resistant, with a high frequency of *kdr* mutations at both loci and high levels of expression of detoxification genes such as CYP6BB2 and CYP6N12. IR03 had a complete knock-down effect against deltamethrin but low mortality and resistance ratios, close to the values for IR05. The IR13 resistance phenotype (96% 24-h mortality) is at the limit of the susceptibility threshold according to the WHO report (24-h mortality > 98%). Our observations of overexpression of CYP6BB2, CYP6N12 and GST2 were in congruence with other publications on French Guiana mosquitoes and another proteomic study using the same strains (Epelboin *et al*., 2021).

On the other hand, our vector competence study revealed less dissemination in the resistant line IR05 than in IR03 and IR13, and we found a negative association between the frequency of 1534C and dissemination saturation level. In a separate study, the risk of infection of *An. gambiae* with *P. falciparum* was positively associated with the L1014F *kdr* genotype in laboratory strains from Senegal and Burkina Faso (Alout *et al*., 2013; Ndiath *et al*., 2014). More recently, the study of Atyame et al. demonstrated that the presence of the homozygous alleles for amplification of the Ester2 locus and homozygous alleles for the acetylcholinesterase ace-1 G119S mutation could enhance the vector competence of *Cx. quinquefasciatus* for West Nile virus (Atyame *et al*., 2019).

Regarding energy storage, the presence of V1016I and F1534C impaired development, longevity and reproduction in *Ae. aegypti* populations in Brazil in the absence of insecticides (Brito *et al*., 2013; Dos Santos *et al*., 2020), because the development of resistance under insecticide-selected pressure is subsequently accompanied by a higher energetic cost or other disadvantages than in susceptible counterparts (Kliot and Ghanim, 2012). In addition, CYP P450-mediated resistance could compete for energy reserves, including lipids and glycogen, in mosquitoes (Hardstone *et al*., 2010). While CHIKV depends on the host’s energetic reserves for its metabolic activities, the energy trade-off caused by resistance could compete with CHIKV development.

Looking at the antiviral capacity of mosquitoes, the existence of a resistant system could simultaneously stimulate their innate immune system by over-expressing detoxification genes, especially the CYP P450 family (Faucon *et al*., 2015). As the CYP 450 are located in oenocytes positioned around the midgut and under cuticles, they can influence the production of reactive oxygen species (ROS) in insects (Murataliev *et al*., 2008). ROS were reported to modulate the immunity of *An. gambiae* to bacteria and plasmodia (Molina-Cruz *et al*., 2008) and were linked to the activation of the Toll pathway for defence against dengue virus infection in *Ae. Aegypti* (Pan *et al*., 2012). Production of ROS in our lines might play a similar role in the different dissemination rates that we observed.

Intriguingly, CYP6BB2, CYP6N12, GST2 were overexpressed for all three lines but trypsin was the only downregulated case, which warrants further study. It was previously reported that some genes coding for serine proteases (including trypsin-like), heat-shock proteins, and C-type lectins were differentially expressed in deltamethrin-resistant populations. Serine proteases are involved in detoxifying insecticide molecules by hydrolysing deltamethrin (Gong *et al*., 2005) and could play an important role in dengue virus infectivity in *Ae. aegypti* (Brackney, Foy and Olson, 2008). The heat-shock protein such as heat-shock protein cognate 70B can help to maintain the homeostasis and inhibit the replication of o’nyong-nyong virus in *Anopheles gambiae* (Sim *et al*., 2007). Taking the *Ae. aegypti* C-type lectin as example, it can be induced by the West Nile virus infection and enhance viral entry by collaborating with a CD45 phosphatase homolog. This positive effect was observed both *in vivo* and *in vitro* studies (Cheng *et al*., 2010). The effects of trypsin on the midgut barrier merit a deeper study.

Focusing back on our experiment, the use of isofemale lines founded from the same breeding site and maintained under intraline breeding had the purpose to attenuate intralineage phenotypic variation and emphasize interlineage ones such as the geographic or adaptive cause of morphological variation (Jirakanjanakit, Leemingsawat and Dujardin, 2008). In addition, this method was used to investigate insecticide resistance conferring pyrethroid resistance in *Anopheles* mosquitoes in Africa (Brogdon *et al*., 1999), and to demonstrate genetic variability in oral susceptibility to arbovirus and vector genotype × virus genotype (G × G) interactions (Lambrechts *et al*., 2009). Nevertheless, even with isofemale lines derived from the same breeding site, the selection of wild mosquitoes based on insecticide pressure could result in lines with different genetic backgrounds. For this reason, in other research, the isogenetic lines which are produced by a series of back-crossings between resistant and susceptible strains are equally used to investigate the interactions between the insecticide resistance and vector competence of *Culex* mosquitoes (Atyame *et al*., 2019; Parker-Crockett *et al*., 2021).

The high infection rates found in all three lines corroborated to previous work by using *Ae. aegypti* populations in South America, including French Guiana, and other CHIKV strains (Vega-Rua *et al*., 2014). The dissemination rates in our experiments were, however, about 50% at 7 DPI, lower than in two other studies, which showed a dissemination rate > 95% at 7 DPI (Vega-Rua *et al*., 2014). The lower vector competence found in this study maybe be due to the following reasons: (1) A lower viral titre was added in the blood meal or the viral clone used is safer but less infectious than the clinically isolated virus; (2) The presence of symbiotic bacteria and/or insect-specific viruses in the midgut of these laboratory lines contributed to natural antiviral defences; (3) Environment factors such as water quality and diurnal temperature range played important roles; (4) The volume of saliva delivered by females which could not be estimated and guaranteed individually, which is a common limitation of studies of vector competence by forced salivation whereas a rapid, simple, specific indicator (marker or colour change reaction indicator) would have been very useful. For example, microRNA is present only in CHIKV-infected mosquito saliva (Maharaj *et al*., 2015), and D7 protein can be detected directly in mosquito saliva by antibodies of human origin (Londono-Renteria *et al*., 2018).

Here we isolated three mosquito lines that differed in both resistance levels and resistance mechanisms and found significantly different midgut barriers among them which affected their vector competence for CHIKV, specifically during the virus dissemination step: the line with the highest resistance had the lowest dissemination rate. These findings might contribute to a better understanding of the interactions between insecticide resistance and vector competence.

## Acknowledgements

We are grateful to Dr Antoine Enfissi for methodological discussions, to Ms Alina Soto for English editing and proofreading the manuscript and to Dr Stanislas Talaga for fruitful discussions.

## Author Contributions

Conceived and designed the experiments: LW, ID, DR, RG, MK

Performed the experiments: LW, PG, AG, JY

Analysed the data: AF, LW, ID

Contributed reagents/materials/analysis tools: MV, AF, DR

Wrote the paper: LW, AF, YE, ID

