## Supplementary materials for "Relevance study of vector competence and insecticide resistance in *Aedes aegypti* laboratory lines"

Lanjiao Wang^1^, Albin Fontaine^2,3^, Pascal Gaborit^1^, Amandine Guidez^1^, Jean Issaly^1^, Romain Girod^1, #a^, Mirdad Kazanji^4^, Dominique Rousset^5^, Marco Vignuzzi^6^, Yanouk Epelboin^1^, Isabelle Dusfour^1^*

### Supporting information

**Fig. S1. Prevalence of infected midguts from IR03, IR05, IR13 and NO and their respective viral loads. A)** Proportions of mosquitoes with an infected midgut over time; 95% confidence intervals are represented by dashes. **B)** Virus load (RNA copies) in midguts over time as determined by qRT-PCR.

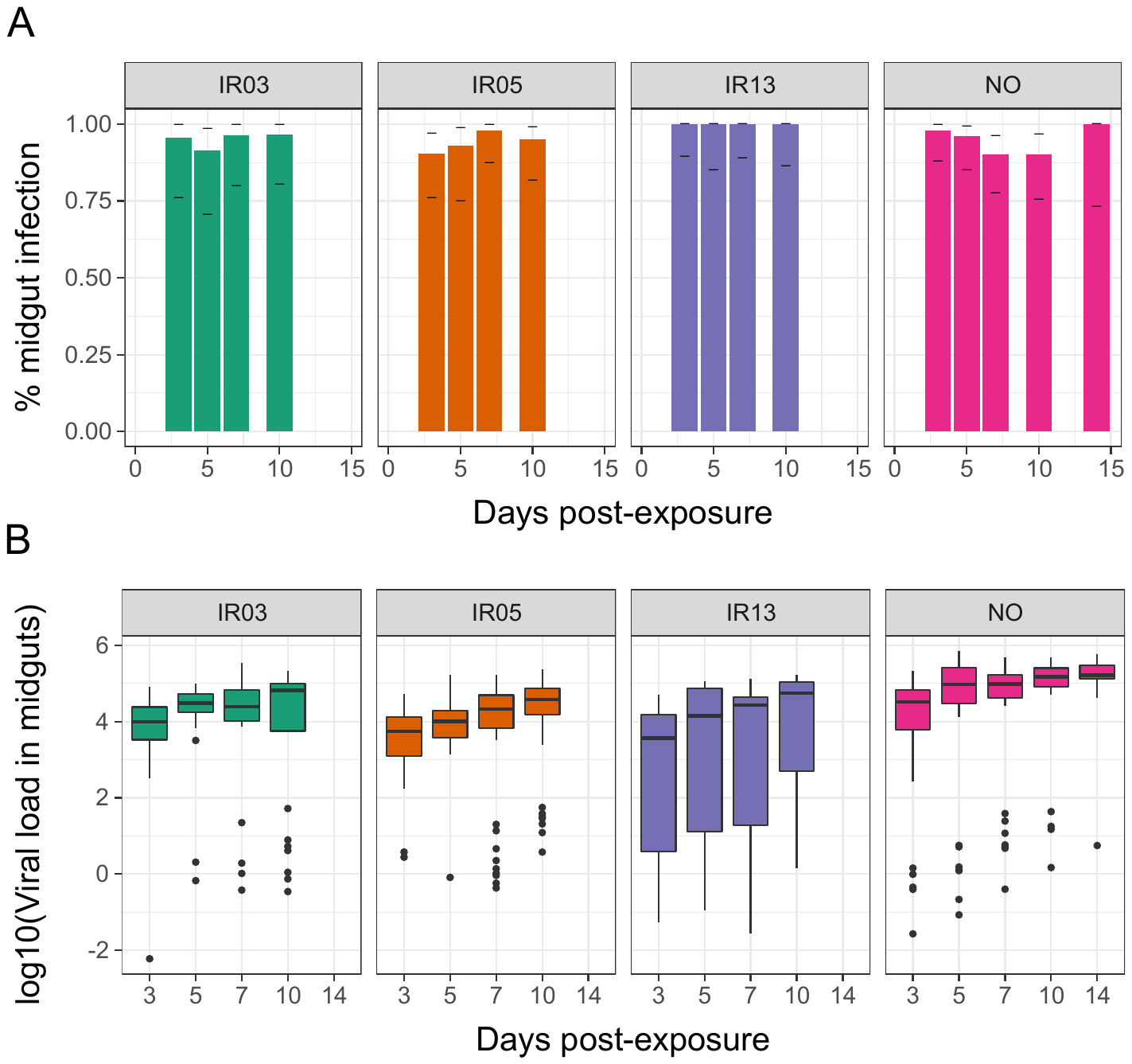

**Fig. S2. Virus load in heads over time as determined by qRT-PCR for each line and the reference NO strain.**

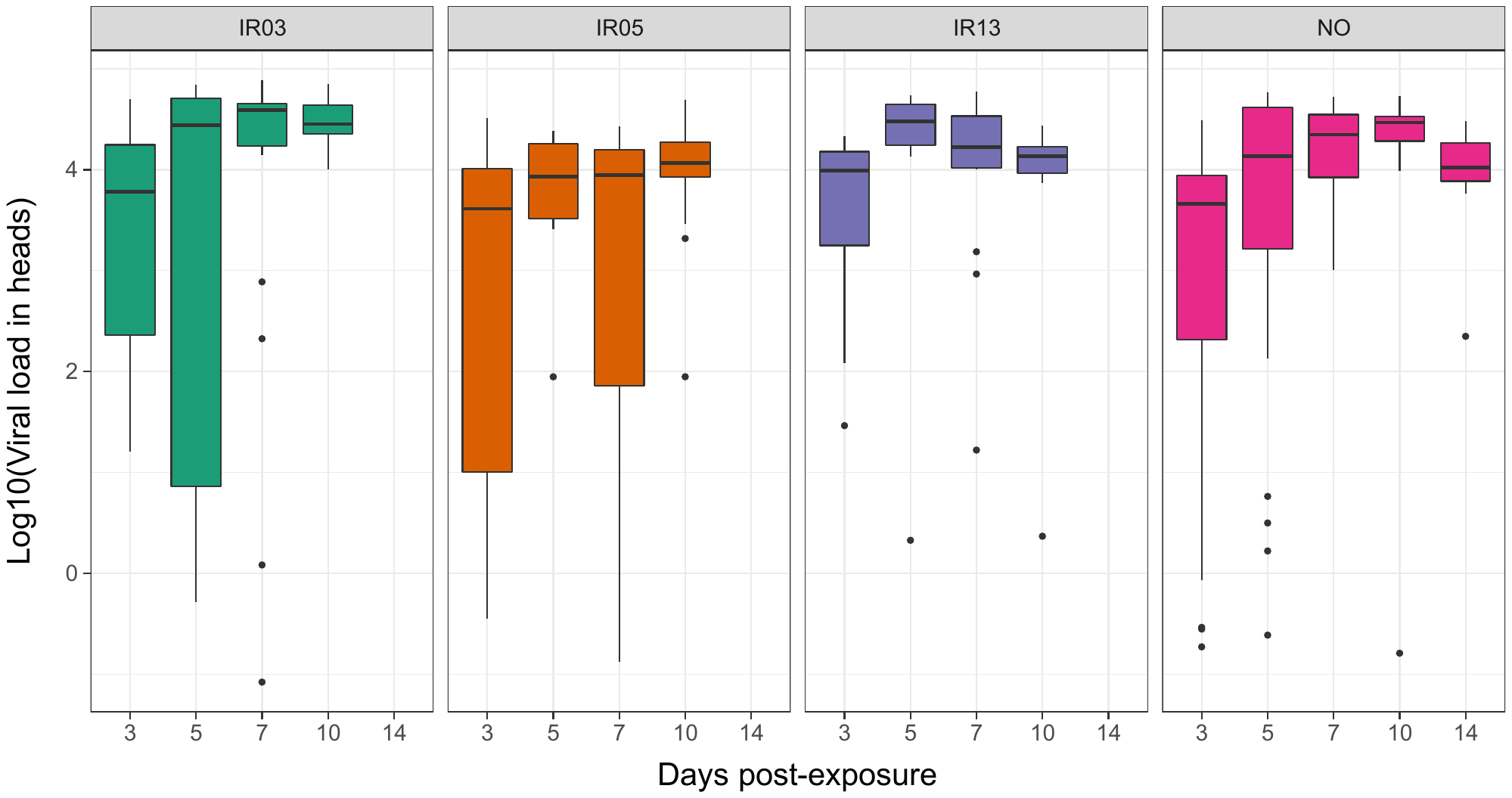

**Fig. S3. Virus load in midgut at 7-day post-infection determined by positive head or negative head for each line and the reference NO strain.**

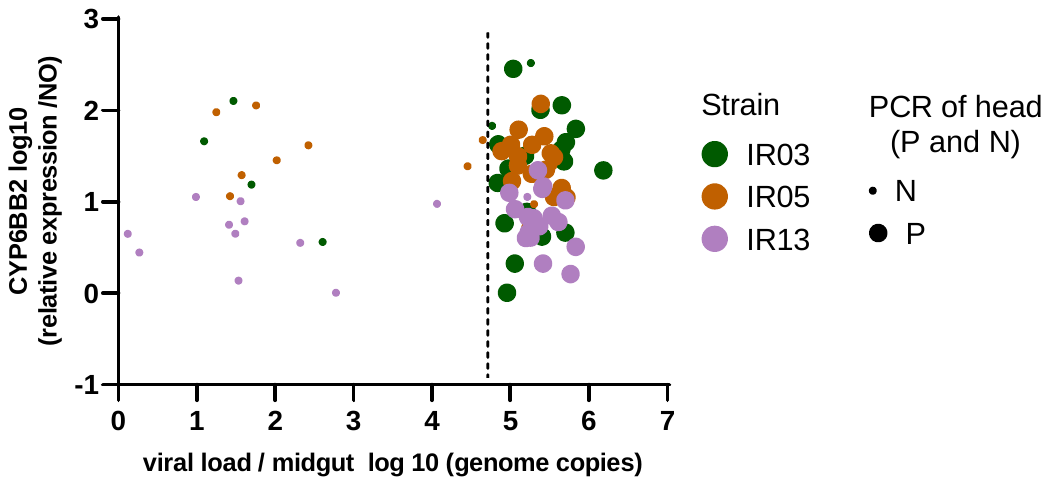

**Table S1. Detailed protocol for qRT-PCR of CHIKV**

| **Name** | **Primer sequence (5'–3')** |
| --- | --- |
| **Primer forward** | TGA-TCC-CGA-CTC-AAC-CAT-CCT |
| **Primer reverse** | GGC-AAA-CGC-AGT-GGT-ACT-TCC-T |
| **Probe** | FAM^1^-TCC-GAC-ATC-ATC-CTC-CTT-GCT-GGC-TAMARA^2^ |
| **Reaction mixture** | 5 µL of RNA template, 5 µL of ddH_2_O, 12.5 µL of RT-PCR buffer (2X), 0.5 µL of 10 µM primer forward, 0.5 µL of 10 µM primer reverse, 0.5 µL of 5 µM probes, and 0.5 µL of Taqman enzyme. |
| **Amplification program** | Reverse transcription at 50 °C for 30 min, activation of Taqman enzyme step at 95 °C for 10 min, 45 cycles at 95 °C for 15 s, 58 °C for 30 s. |

**Table S2. Detailed protocol for kdr genotyping.**

| ***kdr* mutation** | **Primers and probes^1^** | **Primer sequence (5' - 3')** |
| --- | --- | --- |
| **V1016I** | 1016-primer F | GCT-AAC-CGA-CAA-ATT-GTT-TCC-C |
|  | 1016-primer R | CAG-CGA-GGATGA-ACC-GAA-AT |
|  | Val1016-probe | VIC- CAC-AGG-TAC-TTA-ACCTTT-T |
|  | Iso1016-probe | FAM-CAC-AGA-TAC-TTA-ACC-TTT-TC |
| **F1534C** | Phe1534 primer F | GAT-GAT-GAC-ACC-GAT-GAACAG-ATC |
|  | Cys1534 primer R | CGA-GAC-CAA-CAT-CTA-GTA-CCT |
|  | Phe1534-probe | VIC-AAC-GAC-CCG-AAG-ATG-A |
|  | Cys1534-probe | FAM- ACGACC-CGC-AGA-TGA |
| **Reaction mixture** | 3 μL of DNA sample, 12.5 μL of Taqman genotyping Master Mix (Applied Biosystems, USA), 1.8 μL of 10 μM reverse primer, 1.8 μL of 10 μM forward primers, 0.5 μL of 10 μM probe for allele wild-type and 0.5 μL of 10 μM probe for allele mutant, and 0.9 μL of H_2_O. | |
| **Amplification program** | A 95 °C 10-min holding stage, 45 cycles at 95 °C for 15 s, a 60 °C 1-min cycling stage and a 60 °C 1-min post-read stage. | |

**Table S3. Detailed protocol for gene expression.**

| **Name** | **Primer sequence (5' - 3')** |
| --- | --- |
| **RpL8 (**AGAP005802) | Fw-CCT-CGG-GTA-ACT-ACG-CTT-CC  Rv-CCG-CCA-GCA-ACA-ATA-CCA-AC |
| **CYP6BB2 (**AAEL014893) | Fw-AAT-CCC-GAC-ACC-CAT-ACT-GC  Rv-CGC-AGC-AAC-GTA-ACC-AAT-CC |
| **CYP6N12 (**AAEL009124) | Fw-GCG-CTT-CGG-TAT-GAT-GCA-AG  Rv-TCA-CCA-CTC-TAC-AGC-TTC-TGG |
| **GST2 (**AAEL007951**)** | Fw-TGC-CGG-TGC-TAG-ACG-ATA-AC  Rv-ACA-CCG-CTC-TCG-AAG-TGA-AG |
| **Trypsin (**AAEL013712) | Fw-TCC-GAA-ATA-CGA-TGA-TGC-TG  Rv-TAT-TAC-CCC-AGC-CTG-AAA-CC |
| **Reaction mixture** | 7.5 μL SYBR Green Master Mix, 3.5 μL nuclease-free water, 0.5 μL forward and reverse primers (10 μM) and 3 μL cDNA, for a total volume of 15 μL. |
| **Amplification program** | A 95 °C 10-min holding stage, 40 cycles at 95 °C for 15 s, a 62 °C 20-s and a 60 °C 30-s cycling stage and a melt-curve stage. |

**Table S4. Number of positive samples at each collected point.** The table shows the numbers of mosquitoes tested, the numbers of positive midguts, heads and saliva for IR03, IR05, IR13 and NO at 1, 3, 5, 7, 10- and 14-days post-infection. “Infection rate” is the percentage of mosquitoes with positive midguts among total mosquitoes tested; “dissemination rate” is the percentage of mosquitoes with positive heads among all those with positive midguts; “transmission rate” is the percentage of mosquitoes with positive saliva among all those with positive heads; “transmission efficiency” is the percentage of mosquitoes with positive saliva among total mosquitoes tested.

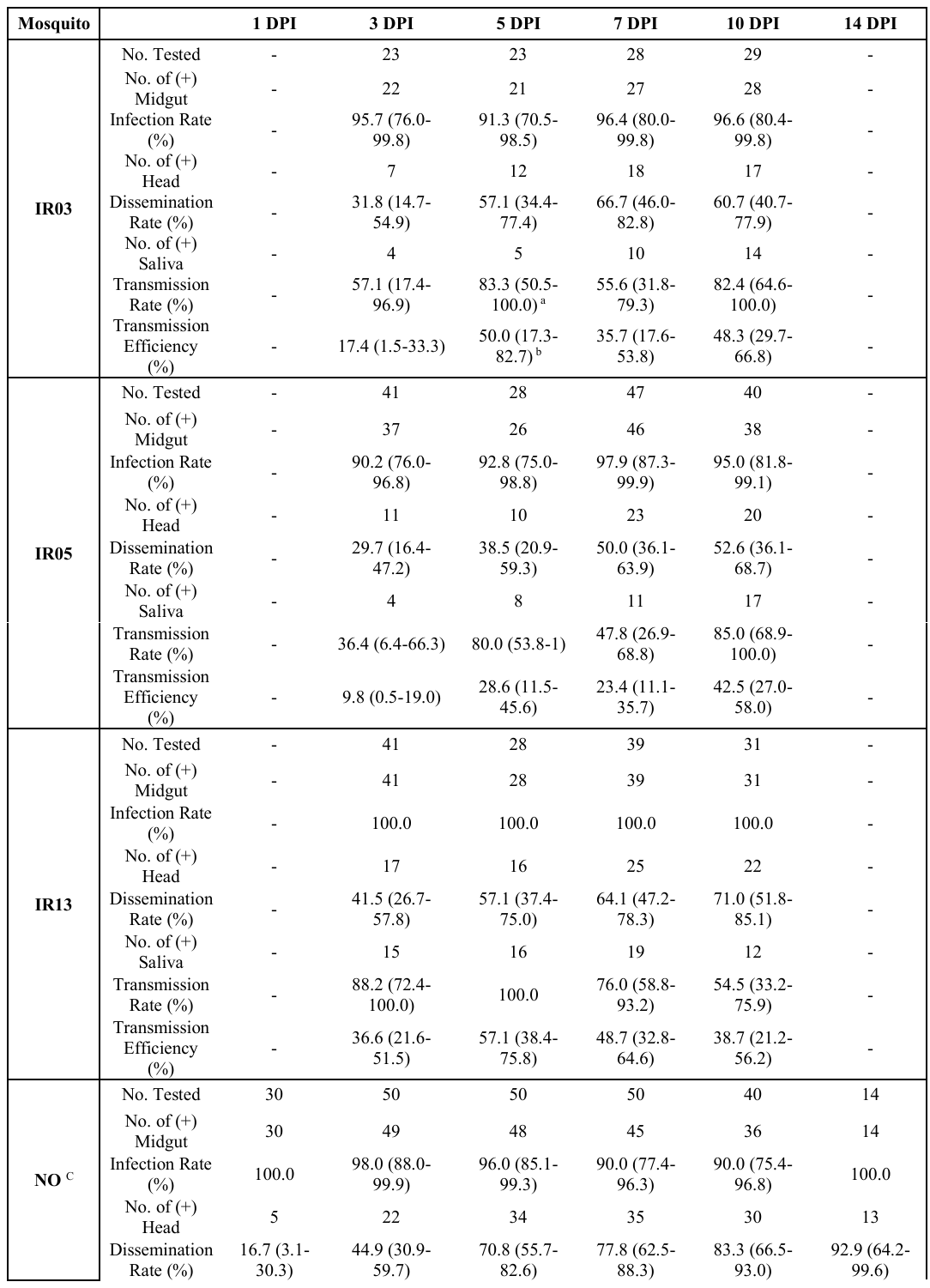
